# Sex-dependent Aβ RA-Low threshold mechanoreceptors alteration in diabetic conditions

**DOI:** 10.64898/2026.10.24.761523

**Authors:** Guillaume Rastoldo, Ali K. Jaafar, Matthieu Bringart, Steeve Bourane

**Affiliations:** Université de La Réunion, INSERM, UMR 1188 Diabète athérothrombose Réunion Océan Indien (DéTROI), Saint-Pierre, La Réunion, France

**Keywords:** DPN, Type 2 diabetes, Mechanical hypoalgesia, Thermal hyperalgesia, LTMR, MafA

## Abstract

Diabetic peripheral neuropathy (DPN) is a major complication of diabetes that profoundly affects quality of life, yet its cellular mechanisms remain poorly defined. Neuropathic pain and sensory loss in DPN are thought to arise from progressive alterations in peripheral nerves usually described as a glove-stocking distribution, but the specific contribution of sensory neuron subtypes has received little attention. Here, we focused on Aβ rapidly adapting low-threshold mechanoreceptors (Aβ RA-LTMRs), a large-diameter sensory neuron subtype best known for mediating touch and vibratory sensation. Using an intersectional genetic approach, we generated Advillin^Cre^; MafA^FlpO^; Ai65 mice, allowing us to selectively label MafA-expressing sensory neuronal populations, which we identified as Aβ RA-LTMRs. We provide the first detailed characterization of how diet-induced type 2 diabetes alters Aβ RA-LTMRs. After 16 weeks of high-fat diet (HFD), we observed striking sex-dependent effects. Male mice developed impaired glucose tolerance, mechanical hypoalgesia, and thermal hyperalgesia, which correlated with a decrease of Aβ RA-LTMR central projections in the dorsal horn. By contrast, female mice showed no changes in either their glycemic status or their sensory sensitivity. These findings demonstrate that Aβ RA-LTMRs are vulnerable to early diabetes-induced alterations in a sex-dependent manner and highlight distinct peripheral versus central mechanisms contributing to sensory dysfunction.

## Introduction

Diabetic peripheral neuropathy (DPN) is one of the earliest and main complications of diabetes. DPN is characterized by neuropathic pain accompanied by nociceptor hyperexcitability, axonal degeneration and loss of cutaneous innervation (Feldman et al., 2019). However, despite the important prevalence of DPN and impact on patients’ quality of life and health care costs, current therapies are only partially effective (Chang and Yang, 2023). Most importantly, the pathophysiological mechanisms leading to neuropathic pain and fiber degeneration have not been elucidated. Studies in neurons and specifically in sensory neurons are necessary to decipher the pathogenesis of DPN. The dorsal spinal cord neurons and circuits that play a crucial role in sensory processing have been the object of little attention in the pathology of DPN (Biessels et al., 2014). Most studies have considered the peripheral sensory neurons has a uniform group (Kobayashi and Zochodne, 2018) but little is known about a potential dysfunction at the level of subtypes of sensory neurons during diabetes.

Somatosensory afferents detect mechanical, thermal, and chemical stimuli in the periphery and transmit these signals to the central nervous system. They are classically categorized into C-, Aδ-, and Aβ-fibers according to their conduction velocities. Their cell bodies are located in dorsal root and trigeminal ganglia, where small-to-medium neurons mainly give rise to C- and Aδ-fibers, while large-diameter neurons predominantly generate Aβ-fibers (Vermeiren et al., 2020). Aβ-afferent neurons remain relatively understudied and are often described as low-threshold mechanoreceptors (LTMR) mediating touch. These neurons give rise to a variety of specialized tactile end organs, including Pacinian corpuscles, Meissner’s corpuscles, and lanceolate endings, which mediate distinct aspects of tactile processing (Handler and Ginty, 2021). DPN affect both small fibers (unmyelinated C fibers and thinly myelinated Aδ fibers) and large myelinated Aβ fibers resulting in sensory and motor deficits (Yang et al., 2025). The recent identification of genetically defined subtypes of sensory neurons and dorsal spinal interneurons (Bourane et al., 2009, 2015; Faure et al., 2020; Sharma et al., 2020; Qi et al., 2024) now provides an opportunity to dissect how diabetes specifically impacts subtypes of sensory neurons. Of particular relevance, the transcription factor MafA is expressed both in Aβ Rapidly Adapting-LTMR (Aβ RA-LTMR) within the dorsal root ganglia, and in a distinct population of dorsal spinal cord interneurons (Bourane et al., 2009). By using intersectional genetic strategy on mice, we aim to selectively target MafA-expressing sensory neuron to investigate Aβ RA-LTMR alterations in a model of diabetic neuropathy.

Here we generated an Advillin^Cre^; MaFa^FlpO^; Ai65 mouse line to provide the first detailed characterization of the impact of early diabetes on Aβ RA-LTMR. We show that high-fat diet (HFD) induces sex-dependent alterations. Male mice develop impaired glucose tolerance, mechanical hypoalgesia, thermal hyperalgesia, and a reduction of Aβ RA-LTMR central projections in the dorsal horn. In contrast, female mice show no detectable impairment in glucose tolerance, sensory sensitivity, or central Aβ RA-LTMR projections after 16 weeks of HFD.

## Methods

### Research Design

All reported experiments were performed at the GIP-CYROI and/or DETROI platform’s animal facility, conducted in accordance with the French and European Community Guidelines for the Use of Animals in Research (86/609/EEC and 2010/63/EU), and were approved by the local Ethics Committee (n°114) for animal experimentation and the French authorities (APAFIS #34213-202108051620151, approved on August 9, 2022).

### Transgenic mouse line

Mice expressing a Cre recombinase under the control of the Advillin promoter (Zurborg et al., 2011) were crossed with MafA^FLpO^ mice (Lecoin et al., 2022) which express FLP recombinase under the control of the MafA promoter, and with Ai65 reporter mice, which carry a double LoxP- and Frt-flanked stop cassette excisable by both Cre and Flp. In Advillin^Cre^; MaFa^FlpO^; Ai65 mice, tdTomato expression is restricted to peripheral sensory neurons that express the MafA transcription factor.

### Experimental design

Twenty-one Advillin^Cre^; MaFa^FlpO^; Ai65 mice (12 males and 9 females) were used in this study. At 8 weeks of age, mice were randomly assigned to one of two dietary groups: a control diet (ND, 10% kcal from fat; SF13-081, Specialty Feeds) or a high-fat diet (HFD, 45% kcal from fat; SF04-001, Specialty Feeds) for 16 weeks. The ND group consisted of n=5 male and n=4 female while the HFD group consisted of n=7 male and n=5 female. Behavioral testing was performed every 4 weeks after the start of the diet and up to 16 weeks. At the end of the 16-weeks dietary intervention, mice underwent an oral glucose tolerance test (OGTT) and nerve conduction velocity (NCV) test prior to tissue collection as previously described (Rastoldo et al., 2026).

### Genotyping

Genotyping of mice was performed by PCR using CreDN2 (GAT CTC CGG TAT TGA AAC TCC AGC), CreUP2 (GCT AAA CAT GCT TCA TCG TCG G), MafA WTs (GAG GCC TTC CGG GGT CAG AGC TTC), MafA WTas (TGTTTC AGT CGG ATG ACC TCC TCC TTG), Ai65mut-Fwd (GCA ACG TGC TGG TTA TTG TG) and Ai65mut-Rv (CCA GCT TGG TTC CCA ATA GA).

### Behavioral Testing

Mice were familiarized with the testing apparatus 1h/day during a week before behavioral testing. Behavioral tests were performed as previously described (Jaafar et al., 2024; Rastoldo et al., 2026). Innocuous haptic touch sensations were measured using brushing assays, thermal sensations were measured by the Hargreaves test and mechanical pain were measure by pinprick and von-Frey assays. Testing was performed every 4 weeks after the start of the diet and up to 16 weeks

### Oral glucose tolerance test (OGTT)

An OGTT was performed after 16 weeks of diet in all mice. After 6 hours of fasting, one drop of tail blood was collected and glycemia was analyzed using a standard glucometer (One Touch Profile, Lifescan Inc., Milpitas, CA, USA, #6 strips). Oral gavage with glucose 30% (2g/kg body weight) into conscious mice was performed and glycemic levels measured 15, 30, 45, 60, 90 and 120-min post gavage.

### Tissue Harvest

After 16 weeks of diet, mice received a subcutaneous injection of Buprenorphine (0.05 mg/kg, Buprecare^®^, Centravet, France) and were anesthetized with isoflurane (3%). Intracardiac perfusion was performed with 20 mL of phosphate buffered saline (PBS) and 20 mL of 4% paraformaldehyde (PFA) dissolved in PBS. The brain, spinal cord with lumbar dorsal root ganglia, sciatic nerve, the fibulae and hind-paw plantar surfaces of each mouse were collected and post-fixed overnight in 4% PFA solution. Biopsies of all tissue collected were then rinsed and cryoprotected into 30% sucrose dissolved in PBS at 4°C. The samples were then embedded in optimal cutting temperature (OCT, Tissue-Tek®, USA) compound and stored at −80°C until sectioning.

### Immunohistochemistry

For immunohistochemistry, the antibodies used were as follows: chicken anti-NFH (Milipore, 1:500), rabbit anti-DsRed (Takara Bio Clontech, 1:500), goat anti-CGRP (Abcam, 1:1000), rabbit anti-PKCγ (Abcam, 1:500) and rabbit anti-S100β (Proteintech, 1:500). Secondary antibody incubations were performed with Alexa Fluor-488, Alexa Fluor-594 and Alexa Fluor-647 conjugated secondary antibodies (1:500). For non peptidergic nociceptor, Alexa Fluor 488-conjugated Griffonia simplicifolia isolectin IB4 (Gs-IB4, 1:300) was used.

### Confocal imaging

Tissues were imaged using a Nikon Eclipse laser scanning confocal microscope equipped with a 40× objective. Glabrous foot skin samples were imaged as Z-stacks with optical sections of 30–50⍰µm, and analyses were performed on orthogonal maximum intensity projections of the original confocal stacks. Sciatic nerve samples were imaged from 14⍰µm-thick tissue sections, whereas spinal cord samples were imaged from 30⍰µm-thick tissue sections.

### Statistical analysis

For behavioral data, a two-way ANOVA for repeated measures followed by Bonferroni’s post-hoc test was used to determine statistically significant differences. For other data, unpaired Student’s *t*-tests (two tailed) were used for statistical comparisons. A *P* value of less 0.05 was considered significant.

## Results

### 1. Genetic targeting of Aβ RA-LTMRs

To study Aβ RA-LTMRs in diabetic conditions, we first generated a transgenic mouse line enabling the specific labeling of these sensory neurons. We used an intersectional genetic strategy (Kim et al., 2009) to specifically label MafA expressing sensory neurons by crossing MafA^FlpO^ mice (Lecoin et al., 2022) with Advillin^Cre^ mice (Zurborg et al., 2011) in which Cre recombinase is expressed in sensory neurons of the DRG throughout the neural axis, including the trigeminal ganglion and hindbrain. The double heterozygous mice were then crossed with a Ai65 reporter mice carrying a dual-stop tdTomato cassette **(Figure 1A)**. This intersectional approach provides high specificity by avoiding tdTomato expression in non-neuronal MafA-expressing cells found in the skin, nerves, and dorsal spinal cord (Bourane et al., 2009).

**Figure 1:**
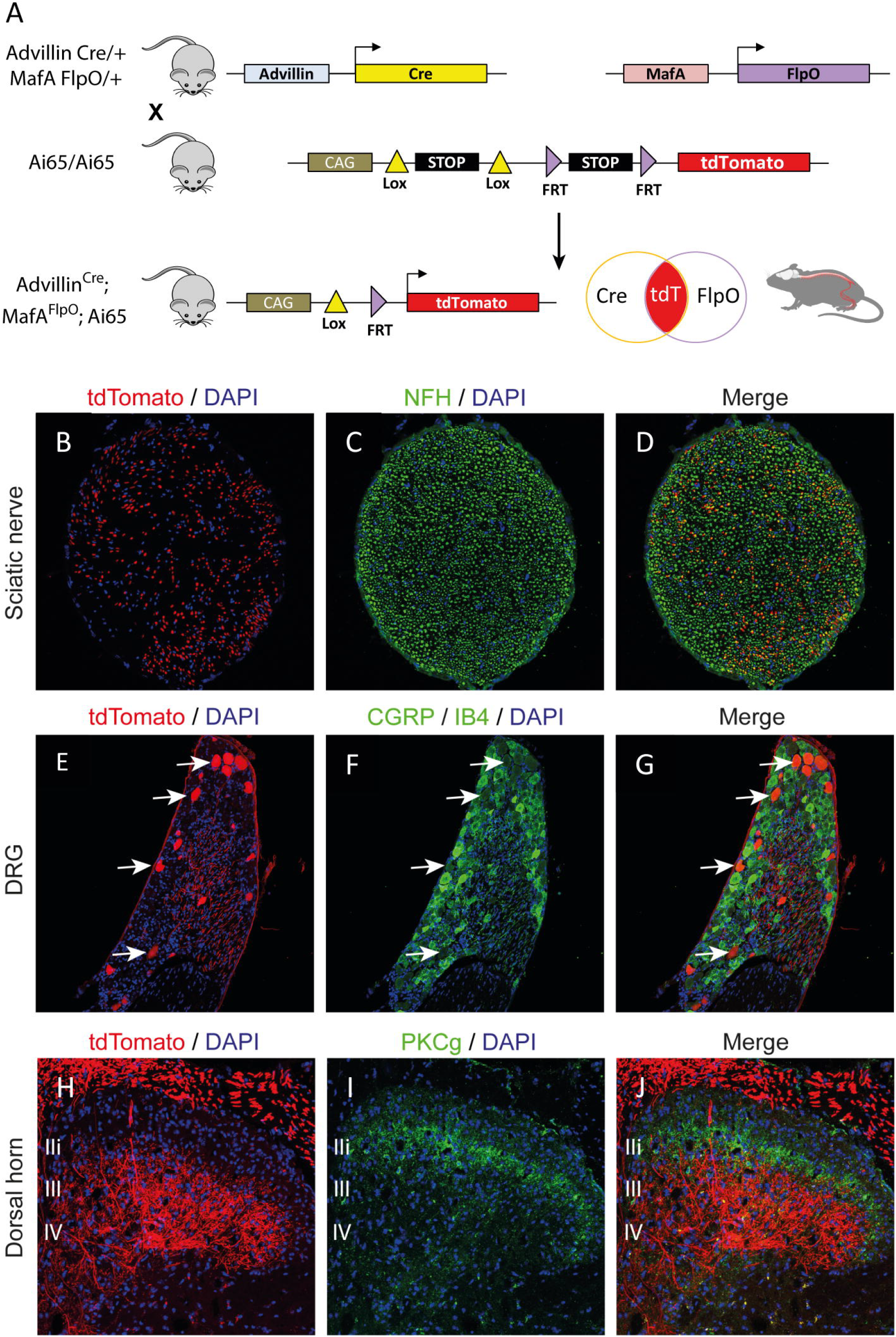
Selective labeling of sensory neurons in Advillin^Cre^; MafA^FlpO^; Ai65. **(A)** AdvillinCre mice, in which Cre recombinase is driven by the Advillin promoter, were crossed with MafAFlpO mice expressing Flp recombinase under the MafA promoter, and with Ai65 reporter mice carrying a dual LoxP- and Frt-flanked stop cassette removable by both Cre and Flp. In the resulting Advillin^Cre^; MafA^FlpO^; Ai65 mice, td-Tomato expression is confined to peripheral sensory neurons expressing the MafA transcription factor. **(B-D)** Immunostaining in the sciatic nerve and DRG **(E-G)** showed specific expression of td-Tomato in subtypes of sensory neurons in Advillin^Cre^; MafA^FlpO^; Ai65 adult mice. **(H-J)** Central tdTomato-positive fibers were predominantly localized to laminae III–IV of the dorsal horn, whereas PKCγ-positive fibers were predominantly localized to lamina IIi.

In the resulting MafA^FlpO^;Advillin^Cre^;Ai65 mice, tdTomato expression was restricted to a specific subtype of sensory neurons in the sciatic nerve **(Figure 1B-D)** and DRG **(Figure 1E-G)** with central tdTomato-positive fibers predominantly localized in laminae III–IV of the dorsal horn **(Figure H-J)**. Peripheral tdTomato-positive sensory endings were observed in lanceolate endings in hairy skin **(Figure 2A-C)**, Meissner corpuscles in glabrous skin **(Figure 2D-F)**, and Pacinian corpuscles in the periosteum of the fibula (Figure 2G-I). These specialized mechanosensory end organs are associated with the detection of skin movement and vibration (Delmas et al., 2011; Roudaut et al., 2012) supporting the identification of the tdTomato-labeled population as Aβ rapidly adapting low-threshold mechanoreceptors (Aβ RA-LTMRs). Additional tdTomato-positive cells with a morphology distinct from sensory nerve endings were occasionally observed in glabrous skin, but were not detected in the sciatic nerve, DRG, or spinal cord (asterisks in **Figure 2D-F**). The identity of these cells was not further characterized.

**Figure 2:**
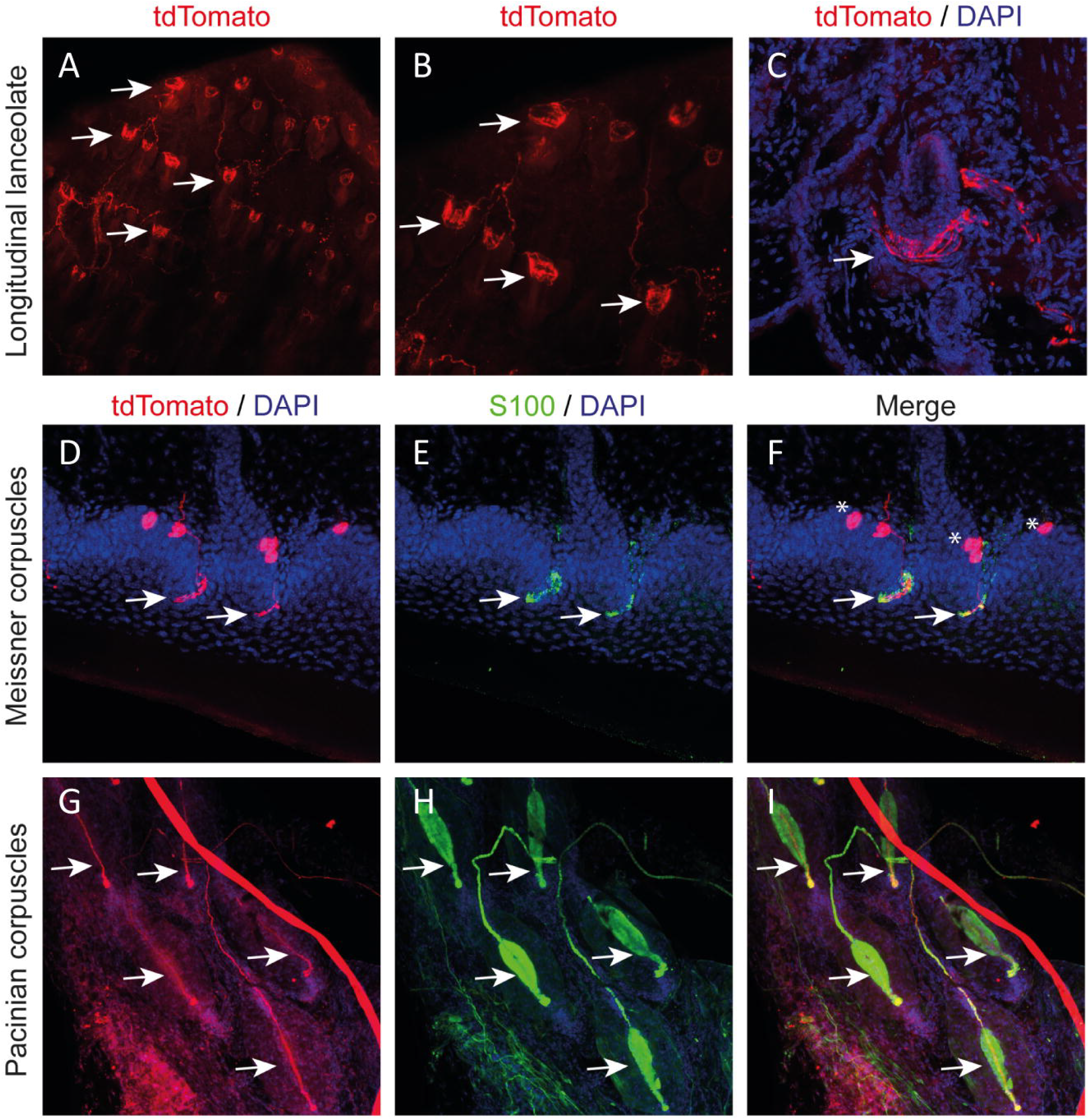
tdTomato labeling identifies Aβ low-threshold mechanoreceptor endings. Sensory nerve endings of tdTomato-labeled neurons in lanceolate endings **(A-C)**, Meissner corpuscle **(D-F)** and Pacinian corpuscle **(G-I)** in Advillin^Cre^; MafA^FlpO^; Ai65 adult mice. White arrows indicate tdTomato-positive sensory endings within the indicated mechanoreceptive structures. Asterisks indicate additional tdTomato-positive cells with a morphology distinct from sensory nerve endings.

### 2. HFD induces sex-dependent metabolic dysfunction with greater impairment in male mice

HFD-fed MafA^FlpO^;Advillin^Cre^;Ai65 mice exhibited significantly greater weight than ND-fed mice after 16 weeks of diet (*P* < 0.001; **Figure 3A**), with male showing a more marked increase (46.70 ± 2.2g; *P* < 0.001; **Figure 3B**) than female (37.81 ± 4.1g; *P* < 0.01; **Figure 3C**). Despite this, fasting glucose levels were not different between HFD- and ND-fed mice (ND: 193.7 ± 17.1; HFD: 223.1 ± 16.4; **Figure 3D**) and no sex-specific differences were observed (male: 227.3 ± 23.9; female: 211.2 ± 21.6; **Figure 3E, F**). Glucose homeostasis was further assessed with OGTT, which revealed a significant impairment in HFD-fed mice, as shown by elevated AUC values (*P* < 0.05; **Figure 3G**). When analyzed by sex, this impairment was present only in males (*P* < 0.05; **Figure 3H**), but not in females **(Figure 3I)**. OGTT curves for total, male, and female groups are shown in **Figure 3J–L**. Our results demonstrate that HFD leads to obesity in both sexes but is associated with sex-dependent metabolic dysfunction, with impaired glucose tolerance observed only in males.

**Figure 3:**
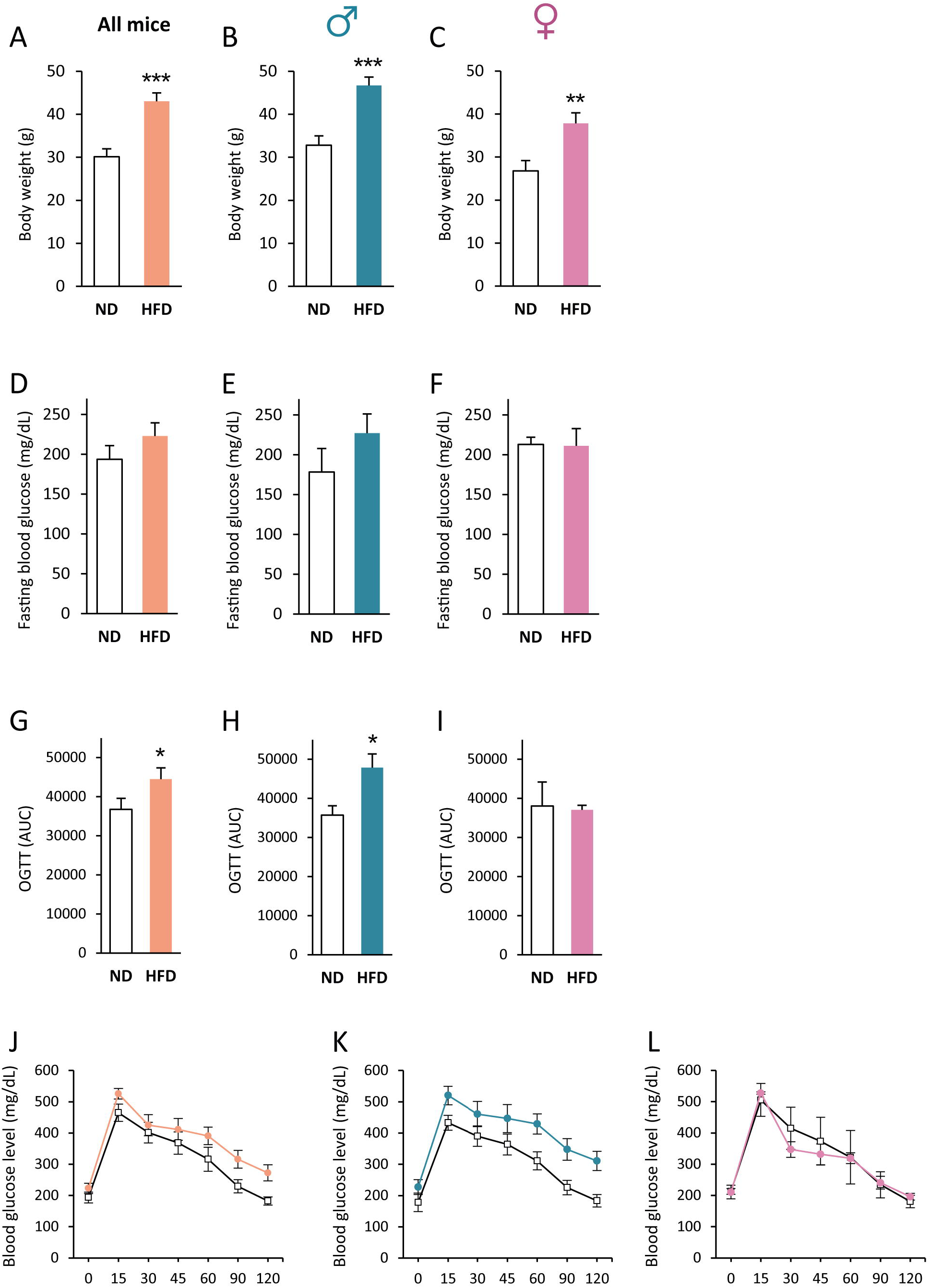
HFD induces greater weight gain and glucose intolerance in male mice. Body weight of ND- and HFD-fed Advillin^Cre^; MafA^FlpO^; Ai65 mice at 16 weeks of diet: total **(A)**, male **(B)**, and female **(C)**. Fasting blood glucose of ND- and HFD-fed Advillin^Cre^; MafA^FlpO^; Ai65 mice at 16 weeks of diet: total **(D)**, male **(E)**, and female **(F)**. Area under the curve (AUC) of Oral glucose tolerance test (OGTT) for ND- and HFD-fed Advillin^Cre^; MafA^FlpO^; Ai65 mice at 16 weeks of diet: total **(G)**, male **(H)**, and female **(I)**. OGTT curves for ND- and HFD-fed Advillin^Cre^; MafA^FlpO^; Ai65 mice at 16 weeks of diet: total **(J)**, male **(K)**, and female **(L)**. Data are presented as means ± SEM.\**p*< 0.05, \*\**p*< 0.01, \*\*\**p*< 0.001: ND vs. HFD.

### 3. HFD induces male-specific alterations in mechanical and thermal nociception

To assess peripheral neuropathy, we performed a series of sensory behavioral tests every 4 weeks over the 16-week diet period. At week 16, HFD-fed mice displayed increased paw withdrawal threshold in the von Frey test, indicating reduced mechanical sensitivity (16 weeks HFD vs. 16 weeks ND, *P* < 0.05, **Figure 4A**). When analyzed by sex, this deficit was specific to male mice (*P* < 0.01; **Figure 4B**), with no significant change in female **(Figure 4C)** suggesting a male-specific mechanical hypoalgesia. Thermal sensitivity emerged in HFD-fed mice at week 8 (*P* < 0.05), which intensified by week 16 (*P* < 0.01; **Figure 4D**). Again, hypersensitivity to heat was specific to male mice (week 8: *P* < 0.01; week 16: *P* < 0.01 **Figure 4E**), with no significant change in female mice **(Figure 4F)** suggesting a sex-dependent susceptibility to diet-induced sensory dysfunction. Sensitivity to acute mechanical pain, measured by the pinprick test, remained unchanged throughout the 16-week dietary intervention, with no significant differences between HFD- and ND-fed mice, regardless of sex **(Figure 5A–C)**. Similarly, innocuous mechanical touch assessed by the brush test showed stable responses across groups and sexes **(Figure 5D–F)**, indicating that these modalities are preserved under HFD conditions. These results reveal sex-dependent sensory dysfunction following HFD, with males exhibiting mechanical hypoalgesia and thermal hypersensitivity, whereas females remain unaffected.

**Figure 4:**
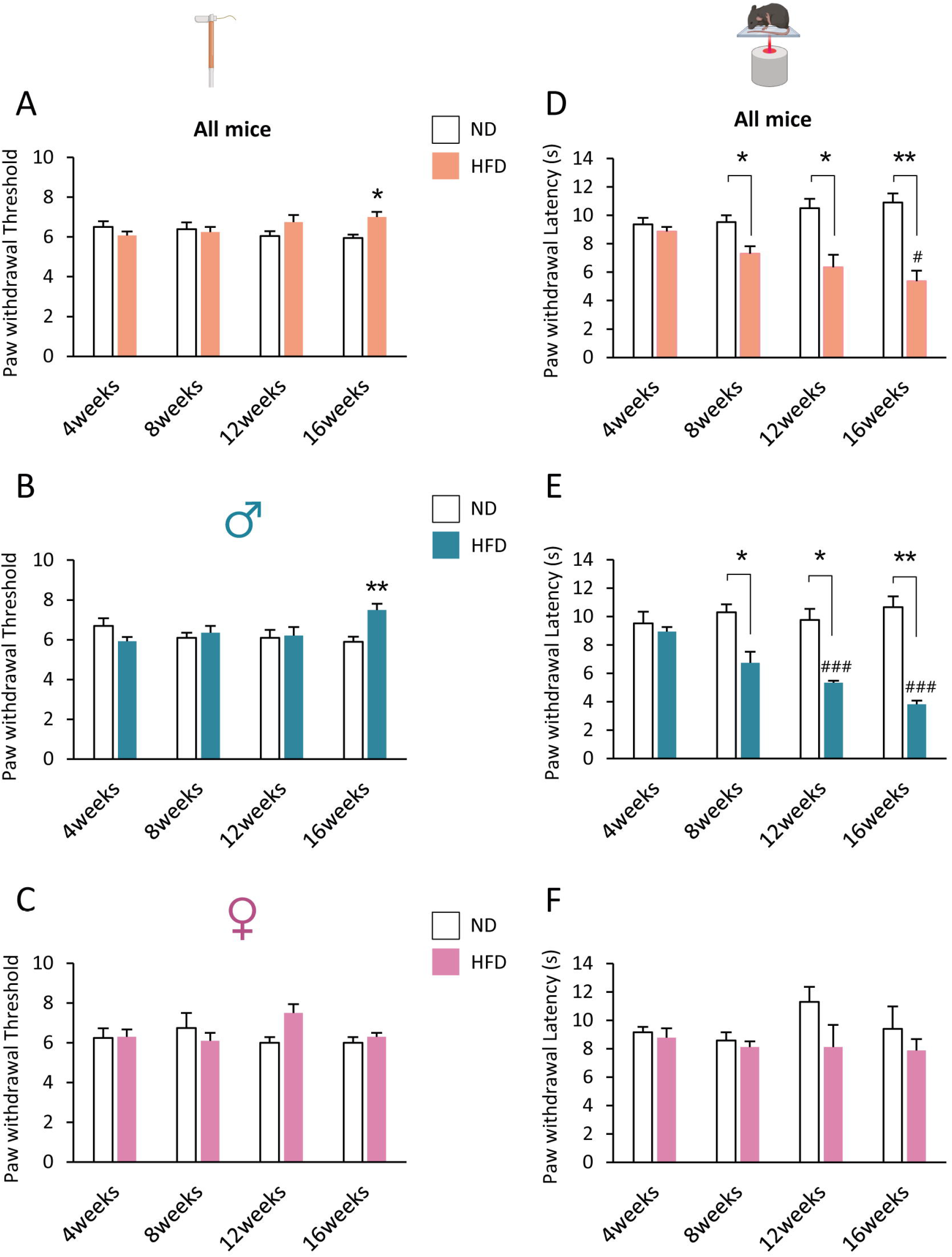
HFD alters mechanical and thermal sensitivity in male mice. Quantification of static mechanical sensitivity by von Frey filaments every 4 weeks in ND- and HFD-fed Advillin^Cre^; MafA^FlpO^; Ai65 mice: total **(A)**, male **(B)**, and female **(C)**. Quantification of thermal sensitivity by Hargreaves test every 4 weeks in ND- and HFD-fed AdvillinCre; MafAFlpO; Ai65 mice: total **(D)**, male **(E)** and female **(F)**. Data are presented as means ±SEM. \**p*< 0.05, \*\**p*< 0.01, \*\*\**p*< 0.001: ND vs. HFD. #*p*< 0.05, ##*p*< 0.01: HFD 4weeks vs. HFD 8, 12 or 16 weeks.

**Figure 5:**
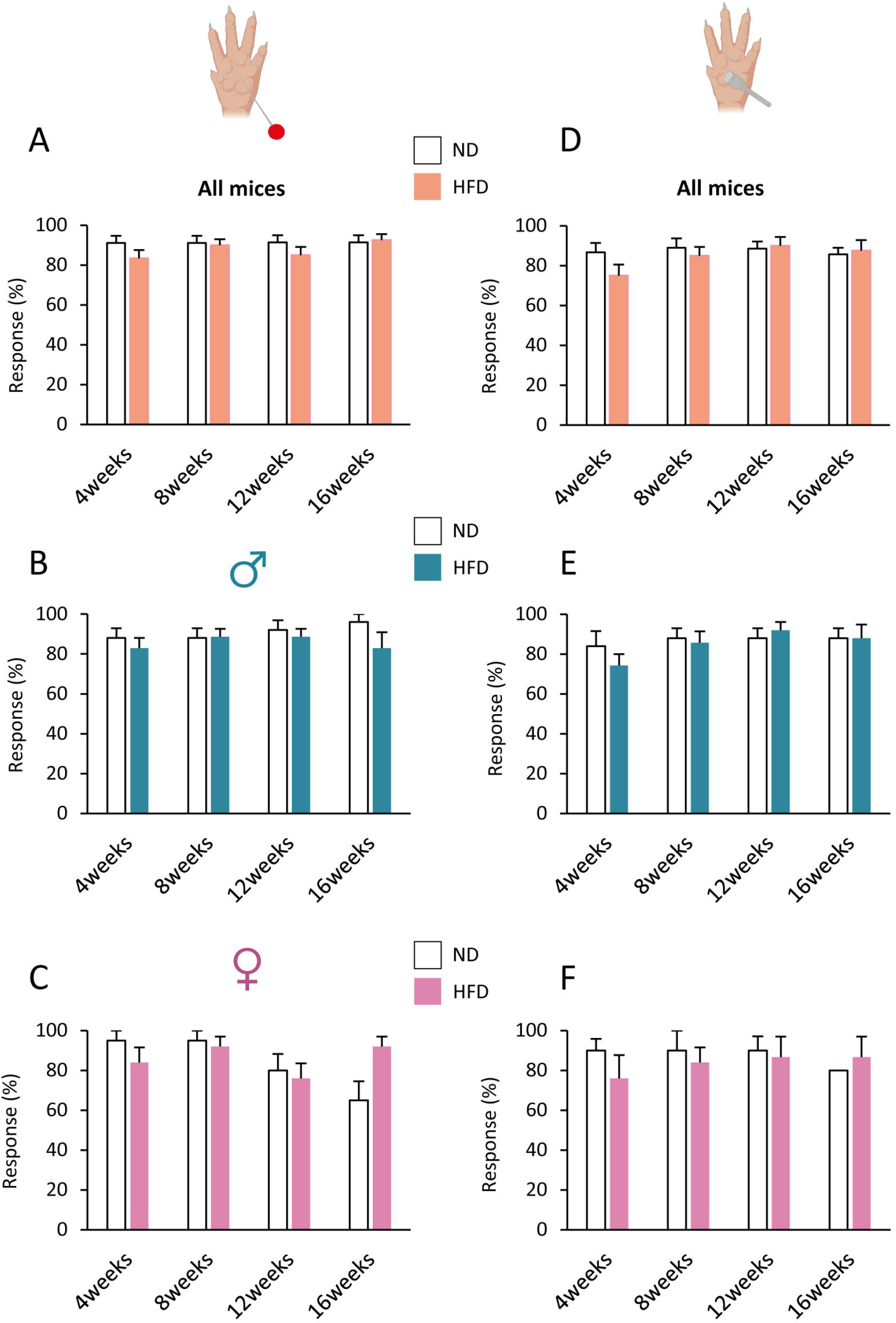
HFD did not alter acute mechanical or innocuous haptic touch sensitivity. Quantification of acute mechanical pain every 4 weeks in ND- and HFD-fed Advillin^Cre^; MafA^FlpO^; Ai65 mice: total **(A)**, male **(B)** and female **(C)**. Quantification of haptic touch sensitivity every 4 weeks in ND- and HFD-fed AdvillinCre; MafAFlpO; Ai65 mice: total **(D)**, male **(E)** and female **(F)**. Data are presented as means ± SEM.

### 4. HFD induces a reduction of tdTomato+ sensory projections in the dorsal horn of male mice

In order to investigate whether HFD-induced sensory dysfunction is accompanied by structural changes in central sensory projections, we quantified tdTomato^+^ labeling in the spinal dorsal horn. We observed that HFD-fed MafA^FlpO^;Advillin^Cre^;Ai65 mice exhibited a significant reduction in tdTomato+ central projection in the dorsal horn of the spinal cord after 16 weeks of diet (*P* < 0.05; **Figure 6A, B**). This reduction is sex-dependent, occurring only in male mice (*P* < 0.001; **Figure 6C**), while female mice exhibited no such change (*P* = 0,3449; **Figure 6D**), suggesting distinct central sensory alterations between sexes under HFD conditions.

**Figure 6:**
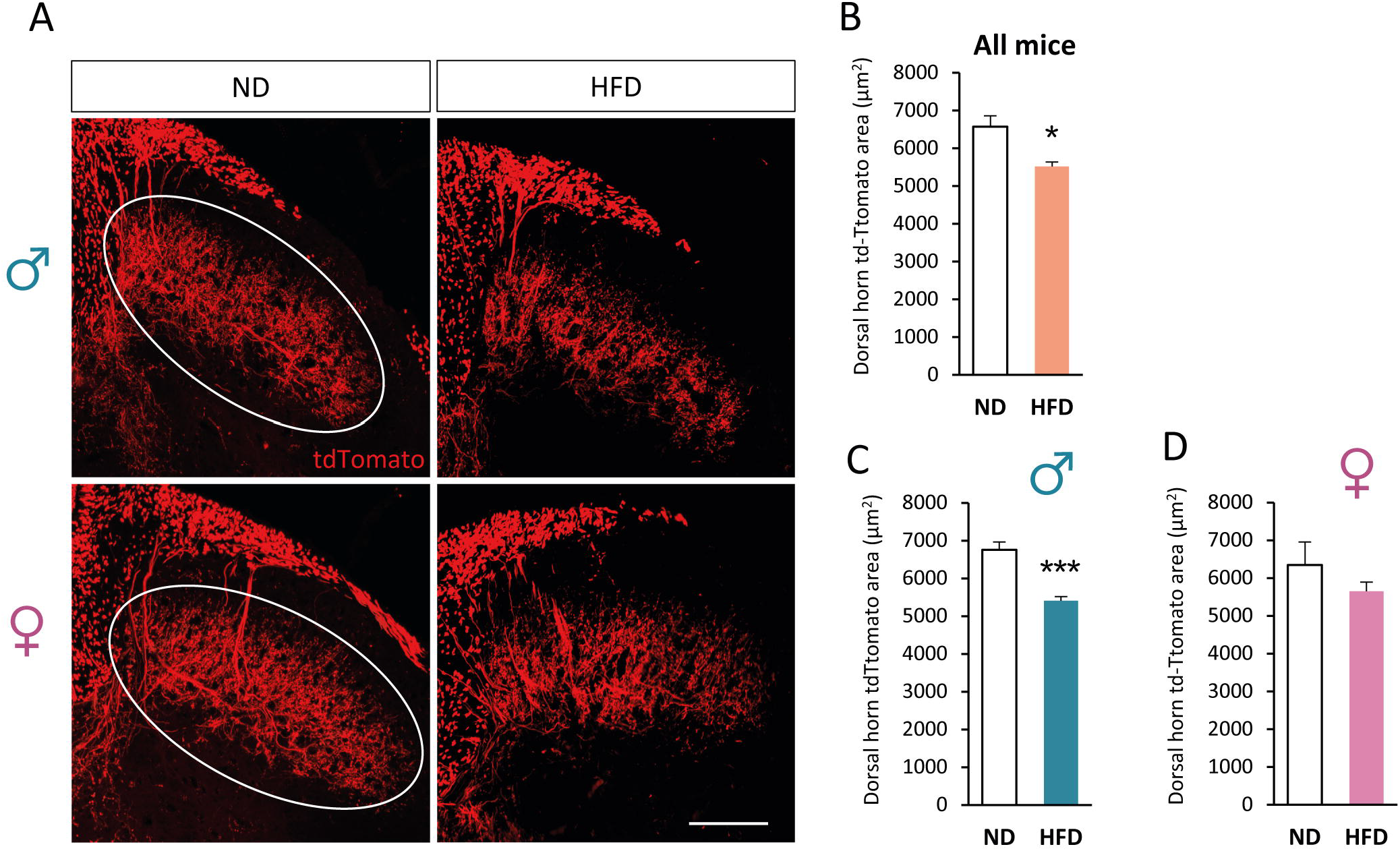
HFD induces loss of tdTomato+ projections in the dorsal spinal cord of male mice. **(A)** Confocal images of dorsal spinal cord cross-sections showing td-Tomato+ (red) projections in male and female Advillin^Cre^; MafA^FlpO^; Ai65 mice fed ND or HFD. Scale bar = 100 µm. Quantification of the td-Tomato+ area in the dorsal horn (white circle) from total **(B)**, male **(C)**, and female mice **(D).** Data are presented as means ± SEM. \**p*< 0.05, \*\*\**p*< 0.001: ND vs. HFD.

## Discussion

Under HFD condition, male mice typically gain weight more rapidly and to a greater extent than female mice. We also observed this effect in our study, with male mice reaching an average body weight of 46.7 g compared to 37.8 g in females, consistent with a relative resistance to diet-induced obesity in females (Daly et al., 2022; Oraha et al., 2022; Saxena et al., 2022; Stapleton et al., 2024). This sexual dimorphism has been attributed to the influence of estrogen, body fat distribution, pro-inflammatory markers, energy expenditure and gut microbiota (Power and Schulkin, 2008; Gaudet et al., 2016; Lainez et al., 2018; Elzinga et al., 2021; Stapleton et al., 2024). Whether these factors also contribute to the divergent neuropathic phenotypes observed in our study remains an open question. Specifically, we observed that male mice developed mechanical hypoalgesia and thermal hyperalgesia, which were correlated with a loss of Aβ RA-LTMR central projections in the dorsal horn of the spinal cord. In contrast, female mice showed no sensory deficits.

Sex-specific pain mechanisms remain poorly understood, in part because decades of research focused primarily on male rodents have produced a male-biased evidence base (Mogil, 2020). Among the biological factors that may contribute to sex differences in pain, sex hormones have received considerable attention (Maurer et al., 2016). For instance, estrogen fluctuations may increase pain sensitivity in females despite their protective immune effects, whereas testosterone exerts analgesic actions by suppressing pro-inflammatory cytokines such as IL-1β and TNFα. In addition, prolactin sensitizes sensory neurons in females, contributing to enhanced pain signaling (Lenert et al., 2021). Despite these insights, the mechanisms underlying sex differences in DPN remain poorly understood, even though painful DPN is reported to be more prevalent in women (Sloan et al., 2018; Elliott et al., 2024). Our findings begin to address this gap by revealing distinct alterations of Aβ-LTMRs between male and female mice. Under physiological conditions, Aβ-LTMR inhibit spinal nociceptive transmission by activating local inhibitory interneurons (Gautam et al., 2024). In male HFD-fed mice, reduced Aβ-LTMR innervation of the dorsal horn would be expected to disinhibit second-order nociceptive neurons, thereby opening the “spinal gate” to pain (Prescott et al., 2014; Gangadharan et al., 2022). Such disinhibition may not only enhance ascending nociceptive signaling but also disrupt top-down modulation of spinal nociception by supraspinal centers (Liu et al., 2018). Surprisingly, our HFD male mice instead showed mechanical hypoalgesia and thermal hyperalgesia. Given the reported role of specific Aβ afferents in gating spinal nociceptive transmission (Sueto et al., 2026), the loss of Aβ-LTMR central projections in HFD-fed males may weaken spinal nociceptive gating that could contribute to their thermal hypersensitivity. In contrast, HFD-fed females exhibited neither sensory deficits nor a loss of Aβ-LTMR central projections.

Our results diverge from those of Elzinga et al., who reported no sex differences in peripheral neuropathy after prolonged exposure to HFD (Elzinga et al., 2021). As discussed in our previous study describing our early diabetes model (Rastoldo et al., 2026), differences in diet composition and mice strain used may help explain this discrepancy. Despite using a transgenic mouse line targeting the transcription factor MafA, we obtained results consistent with those from our earlier work in C57BL/6J wild-type mice (Rastoldo et al., 2026). MafA is best known for its role in pancreatic β-cells, where it regulates insulin gene transcription, glucose sensing, and insulin (Zhang et al., 2005; Artner et al., 2010; Cataldo et al., 2022). However, MafA is also expressed in restricted neuronal populations of the peripheral nervous system, spinal cord, and brainstem (Bourane et al., 2009; Lecoin et al., 2010; Wende et al., 2012; Bikoff et al., 2016; Lecoin et al., 2022). In the present study, we used MafA to identify a subpopulation of mechanosensory neurons, the Aβ RA-LTMRs, and to track their alterations in a mouse model of earlier type 2 diabetes. Interestingly, mutations in the MafA gene have been linked to diabetes mellitus in humans (Iacovazzo et al., 2018). Nevertheless, the insertion of FlpO into the MafA locus in our transgenic model does not appear to predispose mice to diabetes, since animals maintained on a normal diet did not develop prediabetes, in contrast to those fed with HFD.

Diabetic peripheral neuropathy (DPN) remains poorly understood, both in its pathophysiology and natural course (Nawroth et al., 2018). It is generally assumed that the disease begins with microstructural alterations in small unmyelinated C fibers, leading to early deficits in temperature and pain perception, as well as impaired autonomic functions such as sweating and local blood flow. As the disease progresses, loss of C fibers is followed by demyelination and axonal degeneration of larger myelinated fibers (Aβ, Aδ), which is often associated with advanced sensory and motor deficits and may signal the onset of foot ulceration (Feldman et al., 2019; Groener et al., 2020). In our study, however, we observed alterations at an earlier stage: after only 16 weeks of HFD, male mice already exhibited a reduction of Aβ RA-LTMR central projections in the dorsal horn, but not in females. Thus, even in the absence of widespread Aβ fiber degeneration, sex-specific changes in Aβ RA-LTMR plasticity emerge, highlighting central vulnerabilities that may precede the classical progression of DPN.

## Conclusion

Our study provides one of the first characterization of functional subtypes of Aβ RA-LTMRs in the context of diet-induced type 2 diabetes. Our findings reveal sex-dependent susceptibility to HFD-induced metabolic and sensory dysfunction: although both sexes gained weight, only males developed impaired glucose tolerance, altered mechanical and thermal sensitivity, and reduced Aβ-LTMR central projections in the dorsal horn. These structural alterations suggest that Aβ-LTMR remodeling emerges early during diabetic neuropathy and follows distinct sex-dependent trajectories. These findings expand our understanding of how diabetes mellitus affects specific functional sensory neuron subtypes and dorsal horn circuits that process somatosensory information, and how these neurons may be engaged during disease progression. Nevertheless, some limitations should be acknowledged. Our genetic labeling strategy may not capture the entire population of Aβ RA-LTMRs, potentially underestimating the full extent of their alterations. In addition, we did not monitor the estrous cycle in female mice, which could contribute to variability in sensory deficits. Future studies integrating more comprehensive labeling approaches, functional readouts, and control of hormonal fluctuations will be essential to refine these observations and clarify their translational relevance.

## Data availability

Data available on request from the authors.

## Funding

This research was supported by a grant funded by Inserm, Paris, France [ATIP-AVENIR].

## Conflict of interest

The authors report no conflicts of interest in this work.

## Notes

### Competing Interest Statement

The authors have declared no competing interest.

